# Topical TMPRSS2 inhibition prevents SARS-CoV-2 infection in differentiated primary human airway cells

**DOI:** 10.1101/2021.04.23.440619

**Authors:** Wenrui Guo, Linsey M Porter, Thomas WM Crozier, Matthew Coates, Akhilesh Jha, Mikel McKie, James A Nathan, Paul J Lehner, Edward JD Greenwood, Frank McCaughan

## Abstract

**Background:** There are limited effective prophylactic treatments for SARS-CoV-2 infection, and limited early treatment options. Viral cell entry requires spike protein binding to the ACE2 receptor and spike cleavage by TMPRSS2, a cell surface serine protease. Targeting of TMPRSS2 by either androgen blockade or direct inhibition is already in clinical trials in early SARS-CoV-2 infection.

**Methods:** The likely initial cells of SARS-CoV-2 entry are the ciliated cells of the upper airway. We therefore used differentiated primary human airway epithelial cells maintained at the air-liquid interface (ALI) to test the impact of targeting TMPRSS2 on the prevention of SARS-CoV-2 infection.

**Results:** We first modelled the systemic delivery of compounds. Enzalutamide, an oral androgen receptor antagonist, had no impact on SARS-Cov-2 infection. By contrast, camostat mesylate, an orally available serine protease inhibitor, blocked SARS-CoV-2 entry. However, camostat is rapidly metabolised in the circulation *in vivo*, and systemic bioavailability after oral dosing is low. We therefore modelled local airway administration by applying camostat to the apical surface of the differentiated ALI cultures. We demonstrated that a brief exposure to topical camostat is effective at restricting SARS-CoV-2 viral infection.

**Conclusion:** These experiments demonstrate a potential therapeutic role for topical camostat for pre- or post-exposure prophylaxis of SARS-CoV-2, which can now be evaluated in a clinical trial.

## Introduction

A novel subtype of coronavirus, severe acute respiratory syndrome coronavirus 2 (SARS-CoV-2) was first reported in December 2019 and has led to a global pandemic. There is compelling *in vitro* and *in vivo* evidence that angiotensin-converting enzyme-2 (ACE2) and transmembrane serine protease 2 (TMPRSS2) are required for SARS-CoV-2 entry ^1–3^. SARS-CoV-2 binds ACE2 at the cell surface through its spike protein. TMPRSS2 is a serine protease that cleaves the spike protein thus priming membrane fusion and cellular entry ^4^.

A critical question is whether ACE2 and TMPRSS2 are suitable targets for the prevention or treatment of SARS-CoV-2. Any useful intervention targeting viral entry would need to be administered at the early stage of viral infection to target viral replication rather than later phase COVID-19 disease which is thought to be immune-mediated. The main entry point for SARS-CoV-2 infection are the TMPRSS2 and ACE2 expressing ciliated epithelial cells within the nasal cavity and bronchial airways ^5–7^. It follows that for preventative therapies, the use of differentiated airway cells is critical to modelling therapeutic efficacy *in vitro*.

Prior work suggests that soluble ACE2 could act as a decoy receptor for spike protein and reduce SARS-CoV-2 viral load ^8^, and clinical trials testing this strategy are underway ^9^. An alternative approach is to modulate the cleavage of spike protein by inhibiting TMPRSS2 activity, by (1) using androgen receptor (AR) antagonists to suppress androgen-sensitive TMPRSS2 expression ^10 11^, or (2) direct inhibition of the serine protease activity of TMPRSS2 ^2 12^.

TMPRSS2 expression is regulated by androgen signalling via AR binding at 13 and 60 kb upstream of the TMPRSS2 transcriptional start site ^13^. Translocation of this AR binding site is both common and pathogenic in androgen-dependent prostate cancer ^14^. Androgen signalling may underpin the epidemiological evidence linking male gender with worse clinical outcomes in COVID-19 ^15^. Furthermore, a retrospective survey of cancer patients ^16 17^ reported a potential impact of androgen deprivation therapy on COVID-19 severity. As a result, clinical trials of short-course androgen deprivation in early, non-hospitalised male COVID-19 patients have started recruitment (ClinicalTrials.gov Identifier: NCT04446429; ClinicalTrials.gov Identifier: NCT04475601).

Camostat mesylate, a serine protease inhibitor, directly inhibits TMPRSS2 ^18^. Experiments mimicking the systemic delivery of camostat have shown that it is effective in reducing viral entry. Camostat reduced pseudotyped SARS-CoV-2 viral entry in differentiated airway epithelial cells ^2^ and reduced the already modest levels of wild type viral entry in distal airway organoids ^10 19^. A potential benefit of camostat is that it is orally delivered as well as being well-tolerated, and inexpensive ^1^. It has been licensed in Japan since 1985 and is in regular use in Japan and South Korea for patients with chronic pancreatitis. Several clinical trials in patients with COVID-19 are underway (ClinicalTrials.gov Identifier: NCT04455815; ClinicalTrials.gov Identifier: NCT04608266). However, the systemic bioavailability of orally administered camostat is relatively low ^20^.

Given the potential therapeutic role for targeting TMPRSS2 in SARS-CoV-2 infection, we assessed TMPRSS2 regulation by androgens in primary airway cells ^21 11^ and how androgen antagonism or camostat treatment affected SARS-CoV-2 infection in this clinically relevant system. We show that direct inhibition of TMPRSS2 with camostat is effective in reducing SARS-CoV-2 infection, but similar findings were not observed with androgen antagonism. We also demonstrate that topical delivery of camostat at the air-liquid interface effectively reduces SARS-CoV-2 infection in primary airway cells, providing a potentially safe and effective means to deliver TMPRSS2 inhibition to the site of viral entry.

## Methods

### Primary human airway epithelial cell (hAEC) – air-liquid interface (ALI) model

Human airway epithelial cells (hAECs) were purchased from Lonza or were expanded directly from either bronchial brushings from a main airway at bronchoscopy or a nasal brushing from the inferior turbinate from patients at Cambridge University Hospitals NHS Trust (Research Ethics Committee Reference 19/SW/0152). In brief, primary airway cells at passage 2 were expanded in PneumaCult™-Ex Plus Medium (Stemcell; Catalog #05040) then seeded on collagen (Corning; Cat# 354236) coated 24-well transwell inserts with 0.4μm pores (Falcon, Catalog #353095) until fully confluent. Once confluent, the cells were taken to the air-liquid interface (ALI) and cultured with PneumaCult™-ALI Medium (Stemcell; Catalog #05021) for at least 28 days before conducting experiments. On the day of harvest, adherent HBEC cells were washed once with PBS and incubated in Accutase (Stemcell; Catalog #07920) at 37°C for 15 mins. Cells were dissociated by gentle pipetting and then neutralised with DMEM/10% FBS. Whole cell pellets were collected by centrifuge at 300g for 5 min at room temperature. Specific cells used were human bronchial epithelial cells (HBECs) derived from a non-smoking donor (Lonza; Cat# CC-2540, male); human bronchial epithelial cells from a male smoking donor undergoing bronchoscopy for a non-cancer indication, and nasal epithelial cells (MOD006) from a male patient.

Test compounds: 5-dihydroxytestosterone (DHT) (Cat No.S4757 Selleckchem), enzalutamide (Cat No.S1250, Selleckchem), camostat mesylate (Cat. No. 3193 Tocris) were added to the basal chamber of the transwell as indicated and dissolved in DMSO (max final concentration 0.1%) (enzalutamide) or PBS (camostat). Differentiated hAECs were cultured for two days in the absence of androgen stimulation prior to exposure to DHT or enzalutamide for 24 hours. For combination treatments 24 hours enzalutamide pretreatment was applied prior to DHT exposure.

For topical delivery of camostat mesylate, the apical chamber was washed with PBS before 50μl of the indicated concentration of camostat was added to the apical chamber for 15 minutes before being aspirated.

### SARS-CoV-2 infection assays

The SARS-CoV-2 viruses used in this study are the clinical isolates named “SARS-CoV-2/human/Liverpool/REMRQ0001/2020” ^22 23^ and “SARS-CoV-2 England/ATACCC 174/2020 (Lineage B.1.1.7). Stocks were sequenced before use and the consensus matched the expected sequence exactly. Viral titre was determined by 50% tissue culture infectious dose (TCID_50_) in Huh7-ACE2 cells.

For viral infection, the indicated dose was diluted in PBS to a final volume of 50 μL and added to the apical side of transwells containing differentiated HBEC-ALI cultures for 2-3 hours, then removed. At 72 hours post-infection HBEC-ALI transwells were washed once with PBS, dissociated with TrypLE, and fixed in 4% formaldehyde for 15 minutes. Fixed cells were washed and incubated for 15 minutes at room temperature in Perm/Wash buffer (BD #554723). Permeabilised cells were pelleted, stained for 15 minutes at room temperature in 100 μL of sheep anti-SARS-CoV-2 nucleoprotein antibody (MRC-PPU, DA114) at a concentration of 0.7 μg/mL, washed and incubated in 100 μL AF488 donkey anti-sheep (Jackson ImmunoResearch #713-545-147) at a concentration of 2 μg/mL for 15 minutes at room temperature. Stained cells were pelleted and fluorescence staining analysed on a BD Fortessa flow cytometer.

### RT-qPCR

RNA was extracted using RNeasy Mini Kit (Qiagen) according to the manufacturer’s recommendation. Gene expression was quantified using SYBR Green I Dye (Life Technologies) on a QuantStudio 7 Flex Real-Time PCR System (Applied Biosystems). Data was analysed using 2^−◻◻Ct^ method and Design and Analysis Software Version 2.5, QuantStudio 6/7 Pro systems (Applied Biosystems). The following primers were used: ACE2 Forward (5’-3’): CGAAGCCGAAGACCTGTTCTA; Reverse (5’-3’): GGGCAAGTGTGGACTGTTCC; TMPRSS2 Forward (5’-3’): CTGCTGGATTTCCGGGTG; Reverse (5’-3’) TTCTGAGGTCTTCCCTTTCTCCT; TBP Forward (5’-3’): AGTGAAGAACAGTCCAGACTG; Reverse (5’-3’): CCAGGAAATAACTCTGGCTCAT

### Histology

Transwells were washed three times with PBS, then fixed in 10% Neutral buffered formalin. The membrane was removed from the transwell using a scalpel blade and paraffin embedded. 4 μM sections were cut and stained with haematoxylin and eosin.

### Immunofluorescence

Each transwell was washed three times with PBS and fixed using 4% paraformaldehyde (PFA) for 15mins at room temperature before antibody staining. The following antibodies were used: Anti-ACE2 antibody (Anti-ACE2 antibody - N-terminal, ab228349 was originally used. 21115-1-AP, Proteintech was subsequently used - as ab228349 was no longer available; SARS-CoV / SARS-CoV-2 (COVID-19) spike antibody [1A9] (GTX632604, Genetex); anti-SARS-CoV-2 nucleoprotein antibody (MRC-PPU, DA114); Hoechst dye solution (100 μg/ml) was used for nuclei staining. Confocal images were taken using Nikon Confocal Microscopes C2, magnification x40 oil, or entire transwell inserts were imaged using an Arrayscan XTI (Thermo-Fisher) at 10x magnification.

### LDH cytotoxicity assay

For the positive control, 10x Lysis buffer was applied for 45 minutes to the apical surface of HBEC-ALI or nasal-ALI transwells and incubated at 37 C for 45 minutes prior to assay for LDH. For the test samples, the media was removed at the end of the treatment period, then briefly used for washing the apical surface prior to assay. Spontaneous LDH activity was measured by adding 10 μL of sterile, ultrapure water to the apical surface prior to assay. Each condition was prepared in triplicate. Each sample was assessed for LDH release using the CyQUANT™ LDH Cytotoxicity Assay kit (Invitrogen™, C20300) according to the manufacturer’s recommendation. 50 μl of harvested sample medium was added to 50μl of pre-prepared reaction mix and incubated for 30 minutes at room temperature. After stopping the reaction, plate absorbance was measured at 490 nm and 680 nm. % Cytotoxicity was calculated:

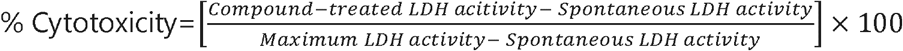

### Quantification and statistical analysis

Statistical analyses of mRNA expression assays and infection quantification data were performed using Prism 8 software (GraphPad Software). Statistical tests used are detailed in figure legends. P values were noted as follows: ns, not significant; *, P < 0.05; **, P < 0.01; ***, P < 0.001.

## Results

### ACE2/TMPRSS2 expression in differentiated human airway cells in response to DHT and enzalutamide

Ciliated airway epithelial cells are the initial site of entry for SARS-CoV-2. To model viral entry in vitro we used differentiated human airway epithelial cells (hAECs) at the air-liquid interface (ALI). Submerged primary hAECs have a basal cell phenotype; they are effectively stem cells of the airway epithelium and are not ciliated ^21 24^. When cultured at ALI in appropriate media, hAECs differentiate to a pseudostratified epithelium that comprise basal, secretory, goblet and ciliated cells ^25^, (**Figure 1A,B**). The expression of ACE2 and TMPRSS2 increased significantly in hAECs cultured at ALI compared to submerged standard culture (**Figure 1C**). This has previously been described for ACE2 (Jia et al., 2005) but these data show that TMPRSS2 is also differentially regulated when airways cells are cultured exposed to air. Given the central importance of both proteins in SARS-CoV-2 infection, these data reinforce the importance of differentiated ALI cultures in studying the initial impact of viral infection.

**Figure 1.**
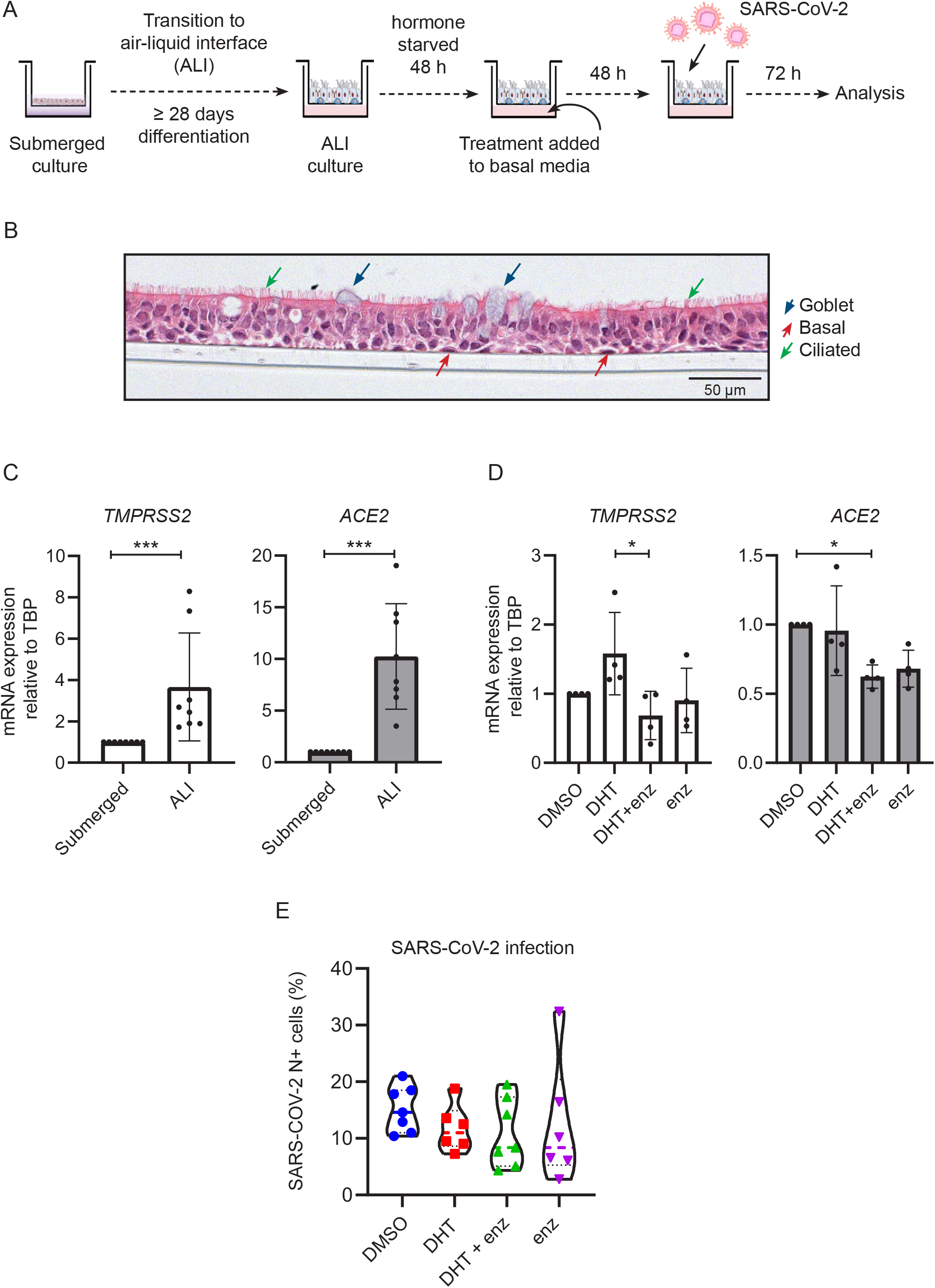
Modulation of androgen signalling does not impair SARS-CoV-2 infection of air-liquid interface differentiated human airway epithelial cells (hAECs). **A.** Schematic representation of the hAEC air liquid interface (ALI) experimental set-up. **B.** H&E staining of fully differentiated pseudostratified epithelium hAEC at the ALI. Ciliated cells, goblet cells, basal cells and are indicated. **C.** hAEC TMPRSS2 and ACE2 expression (mRNA) both increase on culture at the air-liquid interface (ALI) compared to submerged cell culture. p-values were calculated using an unpaired Mann Whitney test. p = 0.0002 and 0.0002 for TMPRSS2 and ACE2 on ALI compared with submerged respectively. **D.** mRNA expression of ACE2 and TMPRSS2 in hAEC-ALI upon treatment with 10 nM DHT with or without 10 μM enzalutamide treatment. p-values were calculated using a Kruskal-Walilis test with Dunn’s correction for multiple comparisons (between all conditions). p = 0.0212 for TMPRSS2 on DHT+enz arm vs. DHT; p = 0.0268 for ACE2 on DHT+enz arm vs. DMSO. **E.** Violin plots showing the quantification of SARS-CoV-2 infected total cells post treating with 10 nM DHT, 10 μM enzalutamide along or in the combination with DHT. All compounds were added to the basal chamber. p-values were calculated using a Kruskal-Walilis test with Dunn’s correction for multiple comparisons (between all conditions). All p-values >0.5.

We next investigated the impact of testosterone signalling on TMPRSS2 and ACE2 expression in our ALI model. Differentiated hAECs were established at the ALI and cultured for two days in the absence of androgen stimulation prior to 24 hours exposure to 5-dihydroxytestosterone (DHT) and/or enzalutamide (**Figure 1A**). DHT is the 5-α-reductase metabolite of testosterone and is a more potent agonist of the androgen receptor than testosterone. Enzalutamide is an androgen signalling antagonist. On treatment with DHT there was a modest, but not significant, induction of TMPRSS2. Treatment with enzalutamide alone had no impact on TMPRSS2 mRNA expression, although it reverted any impact of DHT to baseline. Enzalutamide, in combination with DHT also caused a modest reduction of ACE2 mRNA expression (**Figure 1D**).

### Androgen receptor antagonism does not significantly impact on SARS-CoV-2 infection in human airway epithelial cells

Next, we infected differentiated hAECs with SARS-CoV-2, with or without prior treatment with DHT and/or enzalutamide. These compounds were added to the basal chamber of the transwell mimicking their systemic delivery. 72 hours post inoculation, infection was assessed using a quantitative flow cytometric assay for cells expressing nucleoprotein protein, as a measure of viable infected cells (**Figure 1E**). The well-to-well variation in infection in primary cells has been reported by multiple groups ^2 26 27^. We found no significant impact of either DHT or enzalutamide on the proportion of cells infected in this assay (**Figure 1E**).

### Basal Camostat is effective at blocking SARS-CoV-2 infection

While manipulating androgen receptor signalling was ineffective in modulating SARS-CoV-2 infection, we considered that direct inhibition of TMPRSS2 may have more profound effects. We therefore first treated differentiated hAECs at ALI with camostat for 48 hours in the basal chamber (**Figure 2A**), as a surrogate for the potential impact of systemic therapy, although *in vivo* the compound is metabolised within a minute of reaching the circulation. When delivered in this way, camostat significantly reduced the number of infected cells with or without DHT pre-treatment (**Figure 2B**).

**Figure 2.**
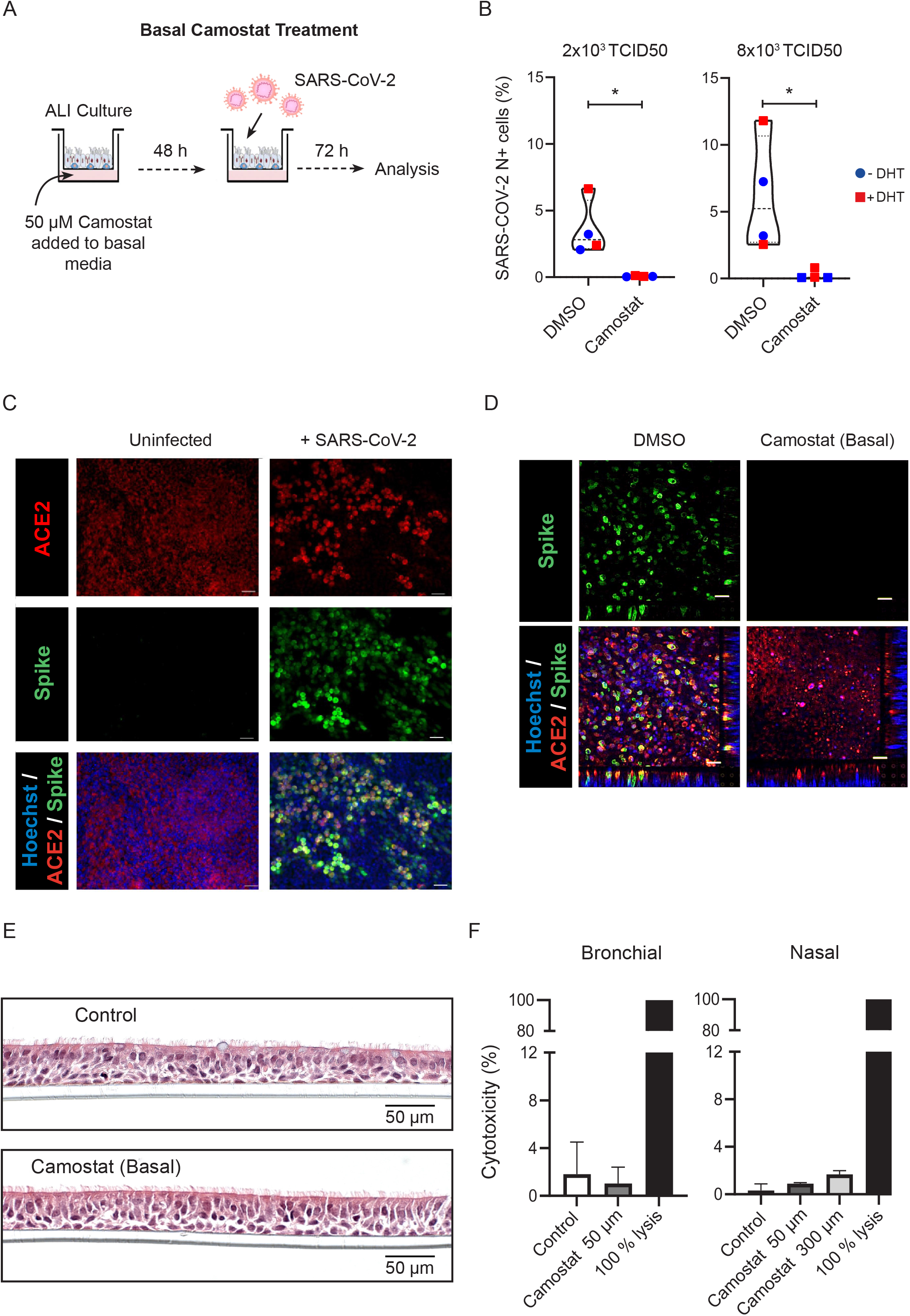
Basal Camostat blocks SARS-CoV-2 infection. **A.** Schematic of experiment. Camostat were added to the basal media 48 h prior to SARS-CoV-2 infection. **B.** Violin plots showing the quantification of SARS-CoV-2 infected total cells after treatment with 50 μM Camostat in the basal chamber with or without DHT. Experiment was repeated twice with wells exposed to either 2×10^3^ or 8×10^3^ TCID50 SARS-CoV2 preparations in 50μl. There was a significant reduction in viral infection in the wells treated with camostat. Samples also treated with DHT are shown in red. p-values were calculated using a Mann-Whitney test, p = 0.0286 at 2×10^3^ TCID50 and p = 0.0286 at 8×10^3^ TCID50. **C.** Representative immunofluorescent image of HBEC-ALI after SARS-CoV-2 infection using antibodies to ACE2 and the S2 subunit of spike protein. Uninfected HBEC-ALI was used to verify specificity. Scale bar:100 μm. **D.** Impact of 48 h of 50 μM basal camostat on cellular infection. Representative immunofluorescent images of hAEC-ALI post SARS-CoV-2 infection using antibodies to ACE2 and the S2 subunit of spike protein. Scale bar: 100 μm. **E.** H&E staining showing normal morphology in cells exposed to 50 μM camostat for 48 h. **F.** LDH release assay as a measure of cytotoxicity following 48 h basal camostat of hAEC-ALI cultures derived from bronchial or nasal cells. Complete lysis shown as a positive control.

Immunofluorescence to a second viral antigen - the S2 subunit of the viral spike protein corroborated the flow cytometry data (**Figure 2C**). Cells were fixed 72 hours after infection. Consistent with previous reports, spike protein colocalised with ACE2 expression. Pre-treatment with camostat markedly reduced the proportion of cells expressing spike protein 3 days after exposure to virus (**Figure 2D**).

Histological analysis of bronchial hAECs, following 48hrs basal camostat treatment showed no toxicity (**Figure 2E**), and there was no excess release of LDH in response to camostat in hAECs-ALI cultures derived from the nose or bronchus (**Figure 2F**).

### Topical camostat retains efficacy in preventing SARS-CoV-2 infection

Having demonstrated that camostat in the basal media effectively inhibits SARS-CoV-2 infection, we hypothesised that topically administered camostat, as a surrogate for nasal or inhaled/nebulised camostat, may prevent SARS-CoV-2 infection. We therefore mimicked local therapy by applying topical 50 μM camostat to the air-exposed side of differentiated hAECs at ALI for either 2 “doses” at 24 hours intervals or a single ‘dose’ approximately 4 hours prior to infection (**Figure 3A**). Each topical dose comprised application of camostat in solution to the apical surface of the hAECs for 15 minutes before removal by aspiration. hAECs were then infected with SARS-CoV-2 and infection analysed at 72 hours by flow cytometry (**Figure 3B**). The proportion of cells infected with SARS-CoV-2 showed a clear dose-dependent reduction. Consistent with the flow cytometry data, immunofluorescence showed that topical treatment with camostat markedly reduced spike protein expression (**Supplementary Figure 1A**). Cross sections of the epithelium stained using nucleoprotein antibody (**Figure 3C**) also corroborated this result. Apical camostat treatment was also effective at reducing SARS-CoV-2 infection in hAEC-ALI cultures derived from nasal cells from an independent donor (**Supplementary 1B**).

**Figure 3.**
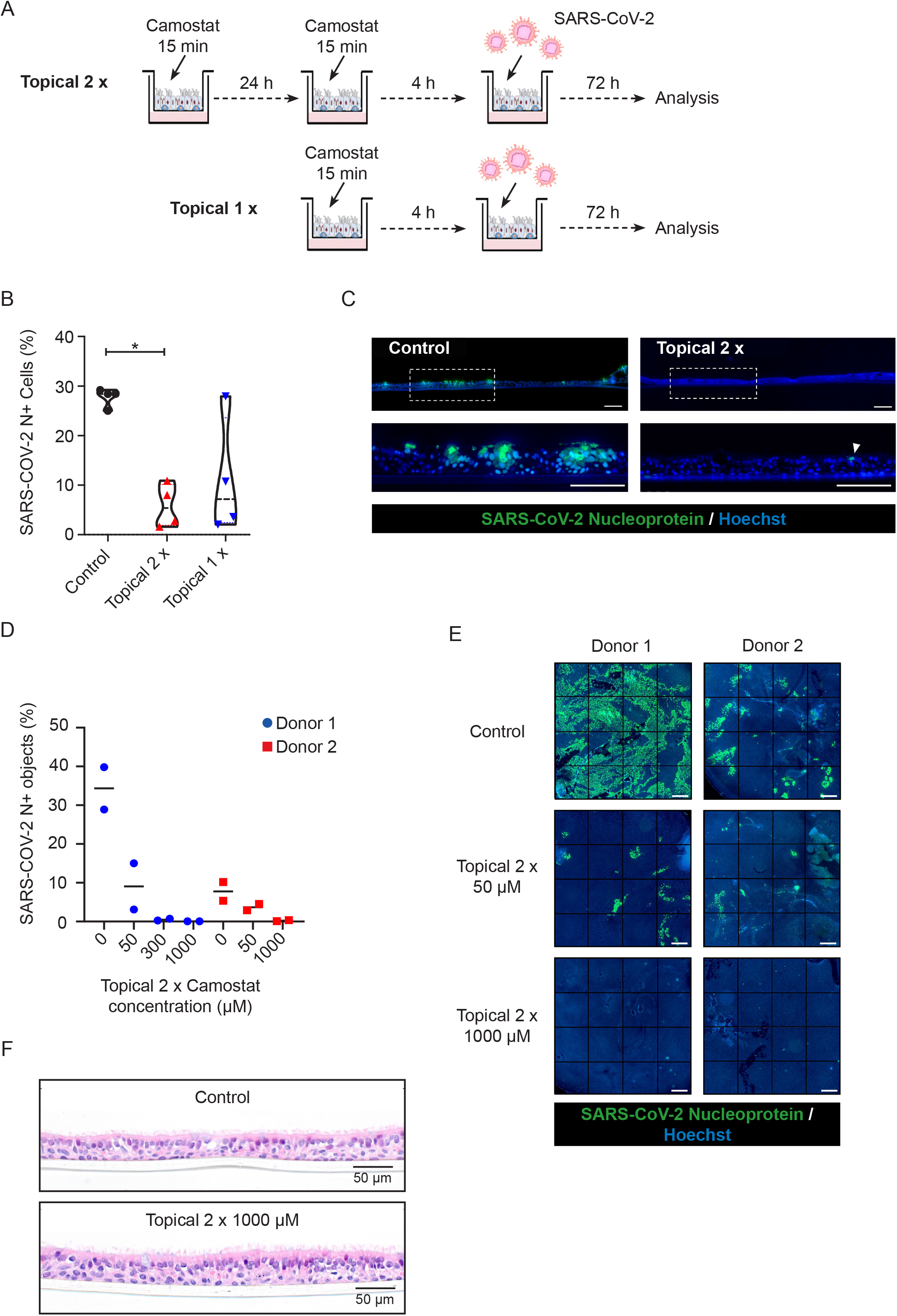
Camostat applied to the apical surface blocks SARS-CoV-2 infection. **A.** Schematic of experimental design for topical camostat treatment prior to SARS-CoV-2 infection. **B.** Violin plots showing the quantification of infected cells when pre-treated with camostat in the apical chamber. p-values were calculated using a Kruskal-Walilis test with Dunn’s correction for multiple comparisons (apical treatment conditions compared with control). p = 0.0285 and p = 0.0997 for topical 2x and topical 1x vs. control, respectively. **C.** Cross-sectional images of IF staining for nucleoprotein on formalin-fixed paraffin-embedded air-liquid interface cultures. Scale bars: 100 μm. **D.** High-throughput microscopy image analysis of hAEC-ALI cultures pre-treated with 2x topical camostat at the indicated dose before infection with 1×10^4^ TCID50 of B.1.1.7 SARS-CoV-2. Bar indicates median. **E.** Example high-throughput microscopy image montages showing the entire scans of representative wells from hAEC-ALI cultures quantified in D. Scale bar: 500 μm **F.** H&E staining showing normal morphology in cells exposed to 2 x topical 1 mM camostat.

Finally, we confirmed that apical camostat treatment was able to block SARS-CoV-2 entry into differentiated bronchial epithelial cells at the ALI derived from a second donor, and using the SARS CoV-2 B.1.1.7 variant. In this experiment, infection was quantified using nucleoprotein staining and high-throughput microscopy. We additionally tested higher dose of camostat, 300μM and 1 mM, again administered for 15 minutes on two occasions separated by 24 hours. The higher dose completely blocked the variant SARS-CoV-2 infection in both donors (**Figure 3D, E**). Topical application of camostat up to 1mM did not induce any gross histological changes in the airway cultures (**Figure 3F**).

## Discussion

### Androgen receptor antagonism is unlikely to be effective at limiting viral entry to airway cells

We have used *in vitro* systems to model the likely impact of androgen antagonism as a means of preventing early SARS-CoV-2 infection ^10 28^. Our experiments suggest that enzalutamide is unlikely to be useful for preventing TMPRSS2-mediated cleavage and subsequent SARS-CoV-2 infection in cells of the conducting airways. A recent commentary questioned the potential use of enzalutamide to prevent COVID-19 ^29^. The authors calculated that 434 individuals would need to be treated with enzalutamide to prevent a single case of COVID-19 and noted the unpleasant side effect profile of enzalutamide ^29^. Our data are in contrast to a recent report suggesting anti-androgens (5-α reductase inhibitors) may reduce SARS-CoV-2 infection in alveolar organoids ^10^. However, there is a respiratory epithelial infectivity gradient with proximal airway cells significantly more susceptible than the distal airway ^30^, and our quantitative assays of airway infection suggest that, compared to camostat mesylate, androgen antagonism is ineffective at preventing proximal airway infection.

### Topical camostat restricts SARS-CoV-2 infection

Our findings show that early administration of topical camostat may be of therapeutic benefit in reducing SARS-CoV-2 infection. Local airway delivery of camostat was previously proposed as an inhibitor of a serine protease that regulates sodium channel flux ^31 32^. The objective was to change mucus characteristics in cystic fibrosis. Coote and colleagues reported preclinical data in hAECs at ALI, in which cells were exposed to up to 30 μM camostat topically for 90 minutes. Camostat inhibition of sodium channel flux persisted at least 6 hours after treatment, despite multiple washes with warmed PBS, demonstrating that topical camostat can have a sustained impact. This duration is consistent with the proposed “pseudo-irreversible” mechanism of serine protease inhibition by camostat mediated by covalent binding to the target ^1^.

A number of *in vivo* preclinical experiments have also been reported, including the administration of nebulised camostat to anaesthetised sheep, ^31^. Up to 60 mg camostat was delivered by nebulisation in 3 ml, equating to a maximal concentration of 50.2 mM, 3 log_10_ higher than we used, without toxicity. Importantly, the duration of activity using a surrogate of therapeutic activity *in vivo*, was at least 5 hours, again consistent with the proposed model of covalent binding to target serine proteases. Further unpublished studies in dogs suggested a mild and reversible inflammatory response in individual animals (lung parenchyma) to inhaled camostat; but no significant toxicity associated with high-dose nasal administration. Therefore, a subsequent early Phase trial focussed on the nasal delivery route - topical camostat was delivered to the nose in volunteers with cystic fibrosis ^32^. They tested dosages in the range 5-1600 μg and used nasal potential difference as a surrogate of target engagement. They reported target engagement and therapeutic efficacy with a 50% maximal effect dose estimated at 18 μg/ml, equivalent to 45.2 μM. Although three serious adverse events were recorded, none were judged to be due to the study medication. Further, the other adverse events were minor and not reported in those administered 200μg or less. Camostat was not detected in the systemic circulation after nasal dosing. The primary metabolite was detected at 3 hours, only in those individuals given the highest dose of 1600μg. Assuming a per nostril epithelial lining fluid volume of 800 μl^33^, the effective concentration after delivery of 1600 μg is 4.0 mM, substantially higher than the doses shown to be effective in this study against two SARS-CoV-2 variants. These data suggest that safe and sustained nasal serine protease inhibition can be achieved using a nasal camostat spray.

TMPRSS2 is a potentially important target for repurposed medications and is an attractive strategy for prophylaxis as the target is the host cell rather than SARS-CoV-2. Both direct inhibition and androgen deprivation have been suggested as strategies to reduce TMPRSS2 activity. Our data does not support the use of androgen receptor antagonists but suggest that direct inhibition of TMPRSS2 will have activity against multiple SARS-CoV-2 variants including B.1.1.7 that has recently been shown to have increased transmissibility ^34^. Consistent with our observations, Hoffmann and colleagues have also recently shown that camostat effectively inhibits variant cell entry in the Caco-2 colorectal carcinoma cell line. They demonstrated this using pseudotyped viral particles engineered to express a series of key variants (B.1.1.7, B1.351 and P.1) implicated as variants that may alter virus-host cell interactions and confer resistance to antibodies ^35^.

Camostat mesylate is generally administered as an oral preparation, but is almost instantaneously metabolised in the circulation to an intermediate metabolite with significantly less activity ^1 36^. Although well tolerated at high oral doses, the maximum concentration of the metabolite attained in the circulation at standard dosing is 0.18μM ^20^, significantly lower than the 10-50 μM concentrations shown to prevent SARS-CoV-2 infection in multiple studies ^1 2 19 20 36 37^.

The impact of an oral camostat preparation in COVID-19 is being actively explored in multiple studies. However, the bioavailability in the airway epithelium after oral delivery remains uncertain. Our data provide a rationale for testing camostat as a local airway administered treatment or prophylaxis for SARS-CoV-2 infection. When effective, local delivery has the advantages of reducing the systemic dose and associated side effects while delivering the drug to the site of disease. Camostat appears well tolerated in the upper airway, a plausibly effective dose was readily achieved and the compound is soluble in saline and cheap to manufacture ^1 31^.

The recent success of the SARS-CoV-2 vaccine trials in protecting against COVID-19 disease should have a major impact on the pandemic ^38–40^. Further, there are now therapeutic options available for treating early phase infection; notably monoclonal antibody administration as single or combination treatments in the pre-hospital setting ^41 42^. This approach is also being tested as a route to pre- or post-exposure prophylaxis in the care home or household contacts setting. The published studies report intravenous administration, although subcutaneous administration is also being trialled.

There remains an urgent unmet need for a safe, easily administered, effective therapeutic with potential efficacy in pre- or post-exposure prophylaxis, and in the reduction of transmission, or progression, in asymptomatic/early SARS-CoV-2 infection. Our work suggests that topical delivery of camostat mesylate should now be clinically evaluated for these indications.

## Acknowledgements

SARS-CoV-2/human/Liverpool/REMRQ0001/2020 was a kind gift from Lance Turtle (University of Liverpool) and David Matthews and Andrew Davidson (University of Bristol). SARS-CoV-2 England/ATACCC 174/2020 was a kind gift from Greg Towers (University College London), and we are also grateful to Ajit Lalvani, Jake Dunning, Maria Zambon and colleagues at Public Health England and Giada Mattiuzzo at the National Institute for Biological Standards and Controls and Wendy Barclay and Jonathan Brown and all colleagues in the United Kingdom Research Institute funded collaboration Genotype to Phenotype. Sheep anti-SARS-CoV-2 nucleoprotein antibody (DA114) was a kind gift from Paul Davies (obtained from MRC PPU Reagents and Services, University of Dundee). LnCAP cells were a kind gift from Charlie Massie. We gratefully acknowledge the support from Dr Ravindra Mahadeva and Ms Jacqui Galloway in establishing the primary cells from patients. We are grateful for the generous support of the UKRI COVID Immunology Consortium, Addenbrooke’s Charitable Trust (15/20A) and the NIHR Cambridge Biomedical Research Centre. This work was supported by a Wellcome Trust Principal Research Fellowship (084957/Z/08/Z) and MRC research grant MR/V011561/1 to P.J.L. This work was supported by the NC3Rs NC/S001204/1 project grant and the Roy Castle Lung Cancer Foundation grant (2015/10/McCaughan) to FM.

This paper presents independent research supported by the NIHR Cambridge BRC. The NIHR Cambridge Biomedical Research Centre (BRC) is a partnership between Cambridge University Hospitals NHS Foundation Trust and the University of Cambridge, funded by the National Institute for Health Research (NIHR). The views expressed are those of the author(s) and not necessarily those of the NIHR or the Department of Health and Social Care.

The authors declare no competing financial interests.

---

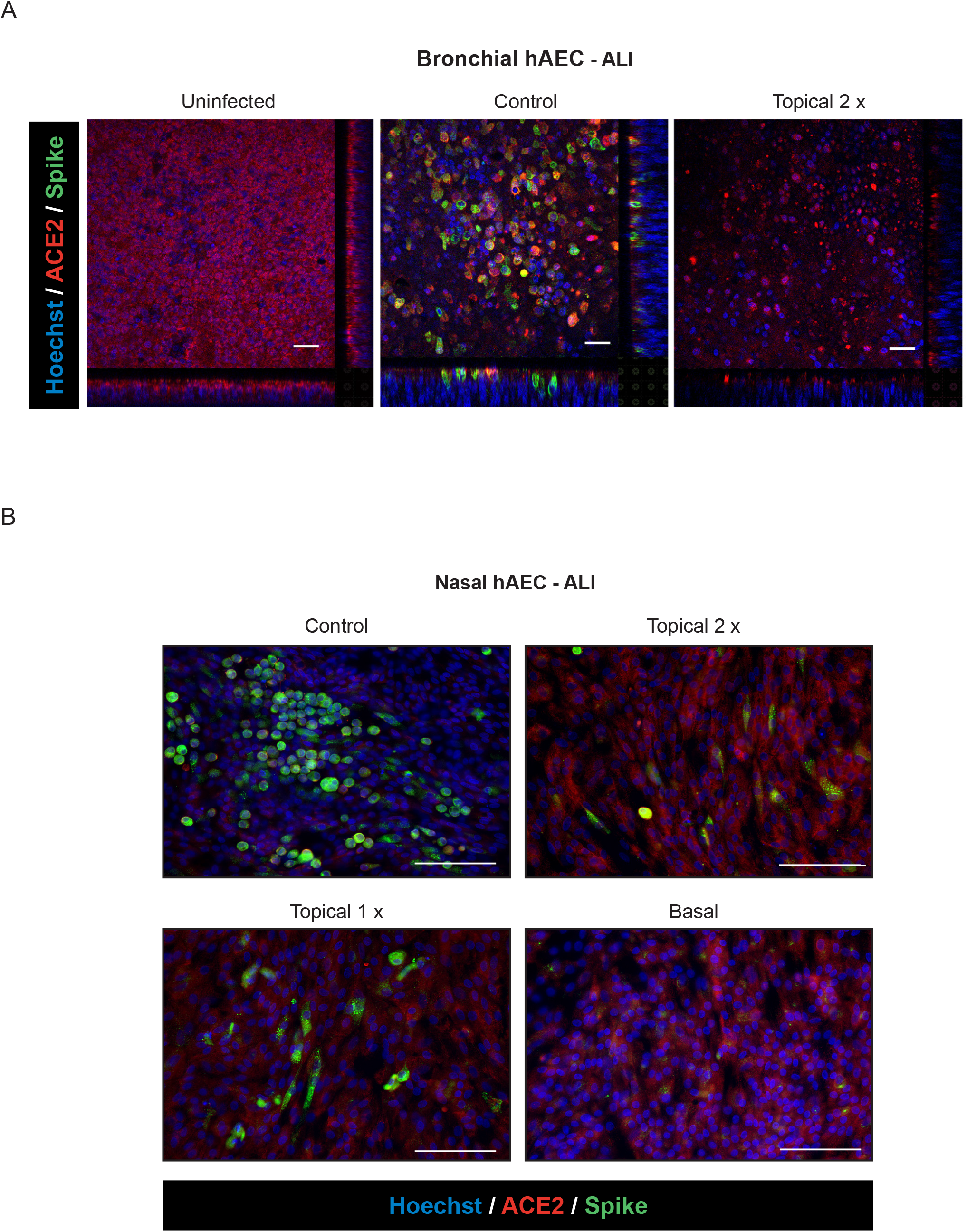

## References

1. Breining P, Frolund AL, Hojen JF, et al. Camostat mesylate against SARS-CoV-2 and COVID-19-Rationale, dosing and safety. Basic Clin Pharmacol Toxicol 2020 doi: 10.1111/bcpt.13533 [published Online First: 2020/11/12]

2. Hoffmann M, Kleine-Weber H, Schroeder S, et al. SARS-CoV-2 Cell Entry Depends on ACE2 and TMPRSS2 and Is Blocked by a Clinically Proven Protease Inhibitor. Cell 2020;181(2):271–80.e8. doi: https://doi.org/10.1016/j.cell.2020.02.052

3. Shang J, Wan Y, Luo C, et al. Cell entry mechanisms of SARS-CoV-2. Proceedings of the National Academy of Sciences 2020;117(21):11727–34. doi: 10.1073/pnas.2003138117

4. Heurich A, Hofmann-Winkler H, Gierer S, et al. TMPRSS2 and ADAM17 cleave ACE2 differentially and only proteolysis by TMPRSS2 augments entry driven by the severe acute respiratory syndrome coronavirus spike protein. J Virol 2014;88(2):1293–307. doi: 10.1128/jvi.02202-13 [published Online First: 2013/11/15]

5. Ruiz García S, Deprez M, Lebrigand K, et al. Novel dynamics of human mucociliary differentiation revealed by single-cell RNA sequencing of nasal epithelial cultures. Development 2019;146(20):dev177428. doi: 10.1242/dev.177428

6. Sungnak W, Huang N, Bécavin C, et al. SARS-CoV-2 Entry Genes Are Most Highly Expressed in Nasal Goblet and Ciliated Cells within Human Airways. ArXiv 2020 [published Online First: 2020/06/19]

7. Martines RB, Ritter JM, Matkovic E, et al. Pathology and Pathogenesis of SARS-CoV-2 Associated with Fatal Coronavirus Disease, United States. Emerg Infect Dis 2020;26(9):2005–15. doi: 10.3201/eid2609.202095 [published Online First: 2020/05/22]

8. Chan KK, Dorosky D, Sharma P, et al. Engineering human ACE2 to optimize binding to the spike protein of SARS coronavirus 2. Science 2020;369(6508):1261–65. doi: 10.1126/science.abc0870 [published Online First: 2020/08/06]

9. Zoufaly A, Poglitsch M, Aberle JH, et al. Human recombinant soluble ACE2 in severe COVID-19. Lancet Respir Med 2020;8(11):1154–58. doi: 10.1016/S2213-2600(20)30418-5 [published Online First: 2020/11/03]

10. Samuel RM, Majd H, Richter MN, et al. Androgen Signaling Regulates SARS-CoV-2 Receptor Levels and Is Associated with Severe COVID-19 Symptoms in Men. Cell Stem Cell 2020;27(6):876–89 e12. doi: 10.1016/j.stem.2020.11.009 [published Online First: 2020/11/25]

11. Ziegler CGK, Allon SJ, Nyquist SK, et al. SARS-CoV-2 Receptor ACE2 Is an Interferon-Stimulated Gene in Human Airway Epithelial Cells and Is Detected in Specific Cell Subsets across Tissues. Cell 2020;181(5):1016–35.e19. doi: 10.1016/j.cell.2020.04.035 [published Online First: 2020/05/16]

12. Stopsack KH, Mucci LA, Antonarakis ES, et al. *TMPRSS2* and COVID-19: Serendipity or Opportunity for Intervention? Cancer Discovery 2020;10(6):779–82. doi: 10.1158/2159-8290.Cd-20-0451

13. Wang Q, Li W, Liu XS, et al. A hierarchical network of transcription factors governs androgen receptor-dependent prostate cancer growth. Mol Cell 2007;27(3):380–92. doi: 10.1016/j.molcel.2007.05.041 [published Online First: 2007/08/07]

14. Rahim S, Uren A. Emergence of ETS transcription factors as diagnostic tools and therapeutic targets in prostate cancer. Am J Transl Res 2013;5(3):254–68. [published Online First: 2013/05/02]

15. Pradhan A, Olsson PE. Sex differences in severity and mortality from COVID-19: are males more vulnerable? Biol Sex Differ 2020;11(1):53. doi: 10.1186/s13293-020-00330-7 [published Online First: 2020/09/20]

16. Montopoli M, Zumerle S, Vettor R, et al. Androgen-deprivation therapies for prostate cancer and risk of infection by SARS-CoV-2: a population-based study (N = 4532). Ann Oncol 2020;31(8):1040–45. doi: 10.1016/j.annonc.2020.04.479 [published Online First: 2020/05/11]

17. Chakravarty D, Nair SS, Hammouda N, et al. Sex differences in SARS-CoV-2 infection rates and the potential link to prostate cancer. Communications Biology 2020;3(1):374. doi: 10.1038/s42003-020-1088-9

18. Shirato K, Kawase M, Matsuyama S. Middle East respiratory syndrome coronavirus infection mediated by the transmembrane serine protease TMPRSS2. J Virol 2013;87(23):12552–61. doi: 10.1128/jvi.01890-13 [published Online First: 2013/09/13]

19. Youk J, Kim T, Evans KV, et al. Three-Dimensional Human Alveolar Stem Cell Culture Models Reveal Infection Response to SARS-CoV-2. Cell Stem Cell 2020;27(6):905–19 e10. doi: 10.1016/j.stem.2020.10.004 [published Online First: 2020/11/04]

20. Bittmann S, Weissenstein A, Villalon G, et al. Simultaneous Treatment of COVID-19 With Serine Protease Inhibitor Camostat and/or Cathepsin L Inhibitor? J Clin Med Res 2020;12(5):320–22. doi: 10.14740/jocmr4161 [published Online First: 2020/06/04]

21. Walters MS, Gomi K, Ashbridge B, et al. Generation of a human airway epithelium derived basal cell line with multipotent differentiation capacity. Respiratory Research 2013;14(1):135. doi: 10.1186/1465-9921-14-135

22. Chu H, Chan JF-W, Yuen TT-T, et al. Comparative tropism, replication kinetics, and cell damage profiling of SARS-CoV-2 and SARS-CoV with implications for clinical manifestations, transmissibility, and laboratory studies of COVID-19: an observational study. The Lancet Microbe 2020;1(1):e14–e23. doi: https://doi.org/10.1016/S2666-5247(20)30004-5

23. Patterson EI, Prince T, Anderson ER, et al. Methods of inactivation of SARS-CoV-2 for downstream biological assays. bioRxiv 2020:2020.05.21.108035. doi: 10.1101/2020.05.21.108035

24. Crystal RG, Randell SH, Engelhardt JF, et al. Airway epithelial cells: current concepts and challenges. Proc Am Thorac Soc 2008;5(7):772–7. doi: 10.1513/pats.200805-041HR [published Online First: 2008/09/02]

25. Sachs LA, Finkbeiner WE, Widdicombe JH. Effects of media on differentiation of cultured human tracheal epithelium. In Vitro Cell Dev Biol Anim 2003;39(1-2):56–62. doi: 10.1290/1543-706X(2003)039<0056:EOMODO>2.0.CO;2 [published Online First: 2003/08/02]

26. Mulay A, Konda B, Garcia G, et al. SARS-CoV-2 infection of primary human lung epithelium for COVID-19 modeling and drug discovery. bioRxiv 2020 doi: 10.1101/2020.06.29.174623 [published Online First: 2020/07/09]

27. Purkayastha A, Sen C, Garcia, G Jr., et al. Direct Exposure to SARS-CoV-2 and Cigarette Smoke Increases Infection Severity and Alters the Stem Cell-Derived Airway Repair Response. Cell Stem Cell 2020;27(6):869–75 e4. doi: 10.1016/j.stem.2020.11.010 [published Online First: 2020/12/02]

28. Bennani NN, Bennani-Baiti IM. Androgen deprivation therapy may constitute a more effective COVID-19 prophylactic than therapeutic strategy. Ann Oncol 2020;31(11):1585–86. doi: 10.1016/j.annonc.2020.08.2095

29. O’Callaghan ME, Jay A, Kichenadasse G, et al. Androgen deprivation therapy in unlikely to be effective for treatment of COVID-19. Ann Oncol 2020;31(12):1780–82. doi: 10.1016/j.annonc.2020.09.014 [published Online First: 2020/10/03]

30. Hou YJ, Okuda K, Edwards CE, et al. SARS-CoV-2 Reverse Genetics Reveals a Variable Infection Gradient in the Respiratory Tract. Cell 2020;182(2):429–46 e14. doi: 10.1016/j.cell.2020.05.042 [published Online First: 2020/06/12]

31. Coote K, Atherton-Watson HC, Sugar R, et al. Camostat attenuates airway epithelial sodium channel function in vivo through the inhibition of a channel-activating protease. J Pharmacol Exp Ther 2009;329(2):764–74. doi: 10.1124/jpet.108.148155 [published Online First: 2009/02/05]

32. Rowe SM, Reeves G, Hathorne H, et al. Reduced sodium transport with nasal administration of the prostasin inhibitor camostat in subjects with cystic fibrosis. Chest 2013;144(1):200–07. doi: 10.1378/chest.12-2431 [published Online First: 2013/02/16]

33. Kaulbach HC, White MV, Igarashi Y, et al. Estimation of nasal epithelial lining fluid using urea as a marker. J Allergy Clin Immunol 1993;92(3):457–65. doi: 10.1016/0091-6749(93)90125-y

34. Davies NG, Abbott S, Barnard RC, et al. Estimated transmissibility and impact of SARS-CoV-2 lineage B.1.1.7 in England. Science 2021;372(6538) doi: 10.1126/science.abg3055

35. Hoffmann M, Arora P, Gross R, et al. SARS-CoV-2 variants B.1.351 and P.1 escape from neutralizing antibodies. Cell 2021 doi: 10.1016/j.cell.2021.03.036

36. Hoffmann M, Hofmann-Winkler H, Smith JC, et al. Camostat mesylate inhibits SARS-CoV-2 activation by TMPRSS2-related proteases and its metabolite GBPA exerts antiviral activity. bioRxiv 2020 doi: 10.1101/2020.08.05.237651 [published Online First: 2020/08/15]

37. Suzuki T, Itoh Y, Sakai Y, et al. Generation of human bronchial organoids for SARS-CoV-2 research. bioRxiv 2020:2020.05.25.115600. doi: 10.1101/2020.05.25.115600

38. Voysey M, Clemens SAC, Madhi SA, et al. Safety and efficacy of the ChAdOx1 nCoV-19 vaccine (AZD1222) against SARS-CoV-2: an interim analysis of four randomised controlled trials in Brazil, South Africa, and the UK. Lancet 2020 doi: 10.1016/S0140-6736(20)32661-1 [published Online First: 2020/12/12]

39. Baden LR, El Sahly HM, Essink B, et al. Efficacy and Safety of the mRNA-1273 SARS-CoV-2 Vaccine. N Engl J Med 2021;384(5):403–16. doi: 10.1056/NEJMoa2035389 [published Online First: 2020/12/31]

40. Polack FP, Thomas SJ, Kitchin N, et al. Safety and Efficacy of the BNT162b2 mRNA Covid-19 Vaccine. N Engl J Med 2020;383(27):2603–15. doi: 10.1056/NEJMoa2034577 [published Online First: 2020/12/11]

41. Chen P, Nirula A, Heller B, et al. SARS-CoV-2 Neutralizing Antibody LY-CoV555 in Outpatients with Covid-19. N Engl J Med 2021;384(3):229–37. doi: 10.1056/NEJMoa2029849 [published Online First: 2020/10/29]

42. Weinreich DM, Sivapalasingam S, Norton T, et al. REGN-COV2, a Neutralizing Antibody Cocktail, in Outpatients with Covid-19. N Engl J Med 2020 doi: 10.1056/NEJMoa2035002 [published Online First: 2020/12/18]

