## Supplementary figure legends for "Topical TMPRSS2 inhibition prevents SARS-CoV-2 infection in differentiated primary human airway cells"

**Supplementary Figure 1. Confocal microscopy showing the effect of camostat treatment on SARS-CoV-2 infection in hAEC-ALI cultures derived from bronchial or nasal cells.**

**A.** Impact of 48 h of topical camostat on cellular infection. Representative immunofluorescent images of bronchial hAEC-ALI epithelia post SARS-CoV-2 infection using antibodies to ACE2 and the S2 subunit of spike protein. Scale bar: 100 µm.

**B.** Impact of 48 h of basal and apical camostat on cellular infection of nasal hAEC-ALI cultures. Nasal ALI epithelial cultures were treated with basal or apical camostat as in **Figures 2A and 3A**. Representative immunofluorescence images of nasal ALI cultures 72 after SARS-CoV-2 infection using antibodies to ACE2 and the S2 subunit of spike protein. Scale bar: 100 µm.
